# Neurobiological slowdown in later life manifests in tempo of popular music

**DOI:** 10.1101/2024.02.06.579086

**Authors:** Geoff Luck

## Abstract

Degradation of motor control across the adult lifespan due to neurobiological decay is well-established. Correspondences between the dynamics of motor behaviour and the timing of musical performance are also well-documented. In light of the former, the conspicuous absence of age as a mediating factor in investigation of the latter reveals a remarkable gap in our understanding of creative performance across the life course. To examine effects of ageing on musical timing, physical tempo of almost 2000 songs released by top-tier recording artists over their decades-long careers were annotated via a listening and tapping task. A series of regression analyses revealed i) an age-driven downward trend in performance tempo for all artists, ii) significant between-artist variation across time, and iii) within-artist variation that was independent of broader musical trends. Overall, tempo decreased by almost one and a half standard deviations from artists’ early twenties to their late fifties, a rate of decline comparable to that observed in studies of spontaneous motor tempo. Results are consistent with the slowing-with-age hypothesis, and reveal that, not only is such tempo decline discernible in commercial recordings, the impact of age on tempo is overwhelming for artists most physically connected with their music.

## Background

The developmental course of human performance in many domains follows a similar pattern across the lifespan. Performance improves throughout childhood into early adulthood, then gradually declines with age. The rate at which one’s abilities taper off appears to be somewhat domain-dependent. For example, a relatively small age-related decline from early-to-mid adulthood onwards has been observed in numerical abilities, verbal ability, and general crystalized knowledge, while a more pronounced degradation has been documented in executive function, memory, processing speed, and reasoning, as well as word production accuracy and recall.^1 2 3 4^ Notably, a weakening of structural and functional connectivity in the brain, especially in sensorimotor systems, and particularly within regions including the primary motor cortex, leads to progressive degradation in motor competence over the same period.^5^

In line with typical lifespan development, motor competence can be conceptualized as a continuous and sequential evolution of motor behaviour that improves from childhood to young adulthood, then degrades steadily across the remainder of one’s life into old age. Consistent with the slowing-with-age hypothesis,^6^ age-related degradation in motor behaviour is thought to reflect an overall slowdown in cognitive, motor, neural, and perceptual processes,^7 8 9^ as well as changes in muscle mass, visuo-proprioceptive function, strength, and reaction time.^10^

Correspondences between the dynamics of motor behaviour and the timing of musical performance are well-documented. For instance, numerous studies have reported connections between biological motion and expressive timing of music.^11 12 13 14 15^ Temporal groupings and lengthening of final phrases similar to those in recorded music have been observed in other physical activities including speech and locomotion.^16 17 18 19^ Cross-cultural studies provide evidence for a common dynamic structure between music and motion.^20^ Music and dance remain inseparable in many cultures.^21 22 23^ A likely explanation for these intimate connections between music and motor behaviour is that throughout the vast majority of human development embodied action has been required to create music.

Conspicuously absent from this work, however, is systematic consideration of age as a mediating factor. This is curious given that the speed of embodied action is known to vary significantly across the life-course.^24^ From early adulthood to old age, performance on speed-dependent motor tasks degrades,^25 26 27^ spontaneous motor tempo (SMT) slows,^28 29 30^ and upper motor rate limit falls.^31^ At the same time, ratings of preferred perceptual tempo are highly correlated with SMT, indicating that individuals also prefer rhythmic stimuli that match those they are able to produce.^32^ Given the universality of music across human cultures^33^ and the well-established degradation of motor timing with age,^34^ such a lack of systematic study reveals a remarkable gap in our understanding of creative performance across the lifespan.

This gap is especially compelling in light of the compositional importance of timing-related features and the powerful impact they exert on listeners’ biobehavioural responses. The most fundamental timing-related characteristic of a musical performance is its underlying speed or tempo. Musicians manipulate tempo to express distinct emotions,^35^ to suggest particular musical styles,^36^ and to build and release tension.^37^ Different tempi influence listeners’ perception of expression,^38^ their level of physiological arousal,^39^ and characteristics of their music-induced body movement.^40^ In light of the well-documented degradation in motor speed across the adult lifespan, the range of tempi an artist produces might logically be expected to slow as they age. This could transform the characteristics and character of their musical output, in turn modifying how listeners engage with their music.

The goal in the present study was to systematically investigate the effect of age on production of musical tempo. To accomplish this, two different approaches were considered: 1) A cross-sectional design in which individuals of different ages are recruited to compose and perform new music, the tempo of which is then compared across age groups; 2) A longitudinal design focused on the tempo of existing music recordings released by culturally-significant artists over their decades-long careers. Compared to the former design, the latter approach has the significant advantages of relatively straightforward acquisition of a much larger dataset, enhanced ecological validity, and considerable relevance to the general population.

Crucially, the longitudinal design also permits within-person comparison of tempo across the same individual’s life course. Indeed, measurement of the same individual across time remains the gold standard in longitudinal studies of lifespan development. Accordingly, a retrospective longitudinal approach was adopted, facilitated by the availability of decades-worth of relevant recordings on music streaming platforms. Specifically, tempi of almost 2000 musical recordings released by top-tier artists over their extended careers were obtained via a listening and tapping task.

Catalogues of the 10 all-time best-selling solo music artists (top 5 female, top 5 male) with careers spanning at least 2 decades were selected for study. Data concerning album sales were obtained from the official Recording Industry Association of America (RIAA) certification database.^41^ A lower limit of 2 decades was applied to increase likelihood of capturing potential age-related decline in artists’ motor speed. According to these criteria, the artists selected were Céline Dion, Elton John, Elvis Presley, Eminem, Lil Wayne, Madonna, Mariah Carey, Michael Jackson, Shania Twain, and Whitney Houston. By nature of their popularity and career length, these artists are arguably the most culturally significant solo artists of the popular music era.

In addition to their general level of popularity and other benefits noted above, there was a further reason for studying such prominent artists. Given their stature in the industry, these artists might be expected to exert significant influence over the characteristics of the music they record, including its tempo. One might thus expect their music to be especially revealing of their tempo production capabilities and characteristics.

Musical tempo itself can be understood as representing two distinct concepts: A physical concept referring to the number of events produced per minute or as a psychological concept representing the event rate perceived by listeners.^42 43^ The majority of scientific work focuses on the latter definition. However, since the goal of the present study was to investigate relationships between event rate created by expert artists (and not that perceived by the average listener) as a function of age, tempo was here defined as the number of events *produced* per minute. To aid comparison between different compositions across different genres and time periods, i.e., to be sure rates of equivalent events were acquired and compared, physical tempo was further specified as the relationship between the kick and snare drums (or equivalents). The rhythmic nature of popular music – the vast majority of which is written in four beats in a bar, with the principal emphasis typically on the first and third beats (kick), and the secondary emphasis on the second and fourth beats (snare) – facilitated this specification. Specifying tempo in this way also helped highlight stylistic trends by removing ambiguity typical in perceptual studies of tempo.

Obtaining physical tempo estimates of large corpora of music is complicated by the overwhelming focus of computational tempo estimation methods on evaluating the *psychological* concept; that is, music information retrieval (MIR) techniques typically attempt to determine the tempo at which an average listener would tap along. Accordingly, the physical tempo of each track was here manually annotated by three musically-trained individuals. Tempi were averaged by album, and relationships between mean album tempo and both year of release and artist age were investigated in a series of regression analyses.

Three hypotheses were formulated and tested. *H1*: Due to physical constraints imposed by a slowdown in motor tempo across the adult lifespan and the intimate connection between biological motion and music performance, artists will exhibit a downward trend in mean album tempo across their careers. *H2*: Artist-specific variations will be observed in these trends due to individual differences in SMT as well as other likely tempo-related influences such as the presence of different producers, musicians, label executives, and genres. *H3*: By nesting tempo within-artist it will be possible to construct a robust overall model in which artist age predicts tempo of musical output.

## Materials and Methods

### Artists

Artists were selected based on two criteria: (i) That they were culturally significant, and (ii) that their careers were of sufficient length to reveal potential age-related effects on tempo. Cultural significance was measured by an index of commercial performance, namely total number of albums sold. Artists often inflate their sales figures to appear (even) more successful than they actually are. To acquire a reliable estimate of performance, sales figures were obtained from the official Recording Industry Association of America (RIAA) certification database. This industry-standard database lists certified album sales for all major artists.

To estimate length of career sufficient to reveal potential age-related decline in musical tempo, literature on human performance across the lifespan was surveyed. Consequently, a meta-analysis was performed on the combined results of multiple studies of SMT by age.^44 45 46 47^ The linear function obtained from this meta-analysis revealed that SMT declines by precisely a quarter (0.25) of a standard deviation per decade across the age range of artists investigated in the present study (18-69). If one assumes that music tempo production more or less mirrors SMT, a minimum career length of 2 decades would be expected to reveal a tempo decrease of approximately half a standard deviation. According to statistical norms, a difference of half a standard deviation is equivalent to a medium effect size. For this reason, when selecting artists, a lower limit of 20 years was placed upon career length. This reduced the likelihood of recent highly successful artists being selected, but increased the likelihood of meaningful age-related effects being identified.

Based on the above criteria, ten culturally-significant recording artists (5 female and 5 male) were selected for study: Céline Dion, Elton John, Elvis Presley, Eminem, Lil Wayne, Madonna, Mariah Carey, Michael Jackson, Shania Twain, and Whitney Houston. According to the RIAA certification database, these artists represent the all-time best-selling solo artists with careers spanning at least 2 decades. Between them, these artists have sold almost 2 billion albums spanning a range of genres including Blues, Chanson, Country, Country pop, Dance, Dance-pop, Disco, Electronica, Funk, Glam rock, Gospel, Hip hop, New jack swing, Pop, Pop rock, Post-disco, R&B, Rock, Rock and roll, Rockabilly, Soft rock and Soul. From a purely statistical viewpoint, an album by one of these artists can be found in every household on the planet.

### Corpus

Initially, the entire back catalogue of each artist was examined. However, given that motor skills are known to improve up to early adulthood and decline thereafter, only songs released after the age of 18 years were included in the current study. Furthermore, to reduce extra-artist influences as much as possible, tempo was not obtained for certain types of albums: Compilation albums, live albums, soundtrack albums, cover albums (those containing 50% or more non-original tracks), Christmas albums, posthumous albums, remix albums, and mixtapes. This produced a corpus of 134 studio albums containing 1866 tracks. From this corpus, several types of tracks were excluded from analysis: Bonus tracks that did not appear on the original release, skits (spoken-word tracks without musical accompaniment), tracks with ambiguous tempi (those for which the tempo was either unclear or varied considerably across time), and tracks with featured artists. The final corpus comprised 1497 tracks spanning 65 years of popular music.

Certified album sales, genres, length of career/album span, and other pertinent information for each artist are shown in Table 1. Album span (number of years from first to most recent album) ranged from 21 (Eminem, Elvis Presley) to 47 (Elton John). Average number of albums per artist was 13, ranging from 5 (Shania Twain) to 30 (Elton John). Average number of tracks per artist was 150, ranging from 55 (Whitney Houston) to 311 (Elton John).

**Table 1.** Certified album sales, genres, length of career/album span, number of albums, and number of tracks for each artist included in the analysis.

| Artist | Certified Album Sales (Millions) | Genres | Album Span (Years) | No. Albums | No. Tracks |
| --- | --- | --- | --- | --- | --- |
| Eminem | 321 | Hip hop | 21 | 10 | 106 |
| Elvis Presley | 231 | Rock and roll, Pop, Rockabilly, Country, Gospel, R&B, Blues | 21 | 21 | 232 |
| Michael Jackson | 248 | Pop, Soul, Funk, R&B, Rock, Disco, Post-disco, Dance-pop, New jack swing | 22 | 6 | 104 |
| Elton John | 204 | Rock, Pop rock, Glam rock, Soft rock | 47 | 30 | 311 |
| Lil Wayne | 165 | Hip hop | 20 | 11 | 120 |
| Mariah Carey | 196 | R&B, Soul, Hip hop, Pop | 28 | 12 | 118 |
| Madonna | 181 | Pop, Electronica, Dance | 36 | 14 | 141 |
| Whitney Houston | 155 | R&B, Pop, Soul, Gospel, Dance | 24 | 6 | 55 |
| Céline Dion | 137 | Pop, Chanson, Soft rock | 32 | 19 | 239 |
| Shania Twain | 81 | Country, Pop, Country pop | 24 | 5 | 71 |
| **TOTAL** | **1 919** |  | **275** | **134** | **1 497** |
| **AVERAGE** | **191,90** |  | **27,50** | **13,40** | **149,70** |
| **SD** | **65,85** |  | **8,62** | **7,92** | **82,92** |

### Annotators

Three individuals (2 female, 1 male, mean age = 24) manually annotated the tempo of each track in the corpus. All were musically-trained students enrolled on a Master’s degree program in musicology at a higher education institution of international repute in the field. Annotators had an average of 12 years of formal instrumental training, played an average of 2 instruments, and had studied music-related disciplines at Bachelor’s and Master’s level for an average of 5 years. These individuals were recruited because of their high-level musical knowledge and demonstrated ability to annotate musical tempo as specified in the present study. Annotators received course credit for their time.

### Data acquisition

Annotators worked remotely, listening individually to assigned playlists of tracks on Spotify. Annotators were unaware of the hypotheses of the study, and each annotator was only debriefed after all had completed their assigned tracks. As a safeguard, tracks were presented in a quasi-randomized order to help mask potential age-related tempo effects. Each annotator tapped the physical tempo of each track using a mobile application and entered it into a database in beats per minute (bpm). Subsequently, each track having a tempo provided by all three annotators was assigned a value in bpm based upon the mean of all tapped tempi. Tracks with tapped tempi deviating from the mean by more than 2 bpm were excluded from analysis.

## Results

### Absolute tempo across all artists

The tempo of musical recordings will likely be affected not only by factors intrinsic to the recordings themselves but also external factors such as changing trends in listener preferences or other market forces. It is important to tease these effects apart in order to identify the most likely source(s) of influence. To this end, year of release (*Year*) and artist age (*Age*) were entered as predictors of absolute mean album tempo (*Absolute Tempo*) in beats per minute (bpm) into a multiple ordinary least squares (OLS) regression. Both *Year, β* = -.85, *t*(131) = -9.43, *p* < .001, and *Age, β* = .53, *t*(131) = 3.62, *p* < .001, were found to be significant predictors of the absolute tempo of commercial recordings of the popular music era, *F*(131) = 45.53, *R*^*2*^ = .41, *p* < .001 (see Figure 1). The large negative effect for *Year* of release suggests that the general tempo of popular music (or at least the music studied here) has gradually slowed since the mid-1950s (Figure 1a). This might reflect changing trends in musical tastes or other market forces across time. However, one drawback of examining tempo in absolute terms is that it obscures between-artist variation: Any differences between artists’ average or how it varies as a function of their age, as predicted by *H2*, could mask the true driver of artist tempo.

**Figure 1.**
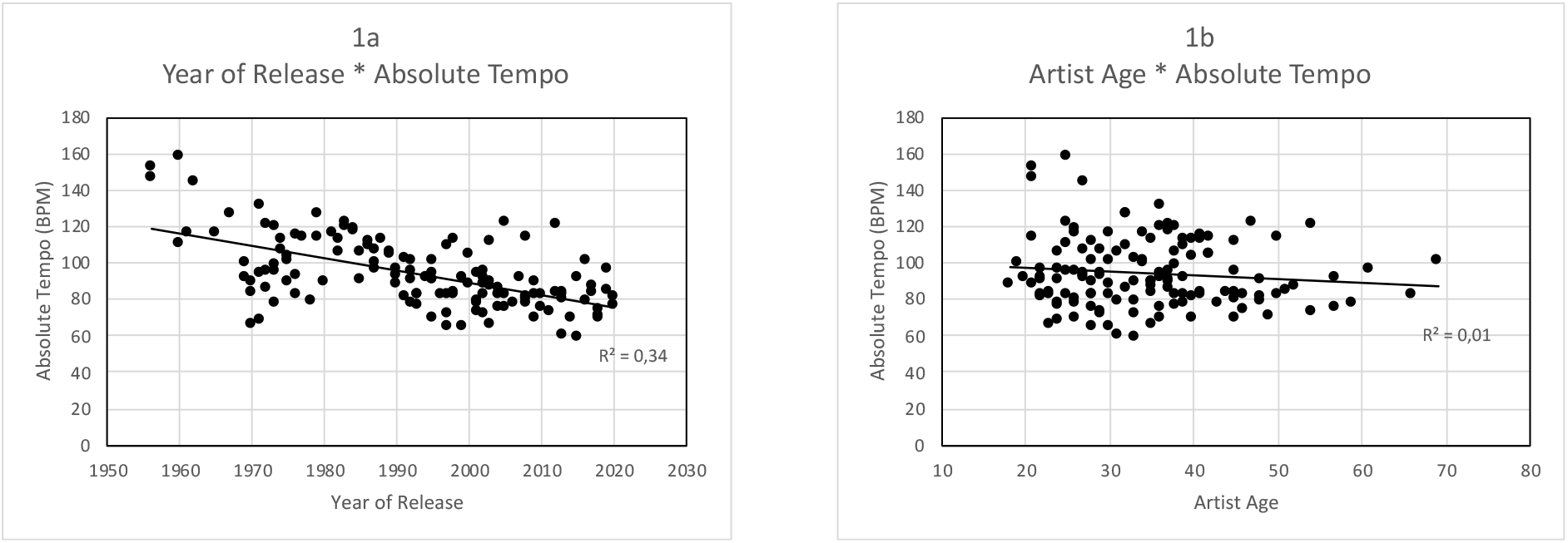
Absolute Tempo as a function of a) Year of Release and b) Artist Age.

### Standardized tempo across all artists

Consequently, *Absolute Tempo* was standardized (z scored) within each artist to create *Standardized Tempo*. This allowed direct comparison between artists’ tempo as a function of age. In a second multiple OLS regression, *Year* and *Age* were entered as predictors of *Standardized Tempo*. The overall model was significant, *F*(131) = 8.72, *R*^*2*^ = .12, *p* < .001. Notably, while *Age* had a significant negative effect on *Standardized Tempo, β* = -.03, *t*(131) = -3.28, *p* = .001, the effect of *Year* was not significant, *β* = .00, *t*(131) = -.51, *ns* (see Figure 2). This suggests that within-artist tempo variation across time is unaffected by broader musical trends. Instead, such variation is at least partly driven by artists’ advancing age. Analysing all artists together, however, even after standardizing tempo, still leaves potential for between-artist variation to obscure possible age-related effects. This finding nonetheless offers support for *H1* and motivates investigation of tempo as a function of age separately for each artist.

**Figure 2.**
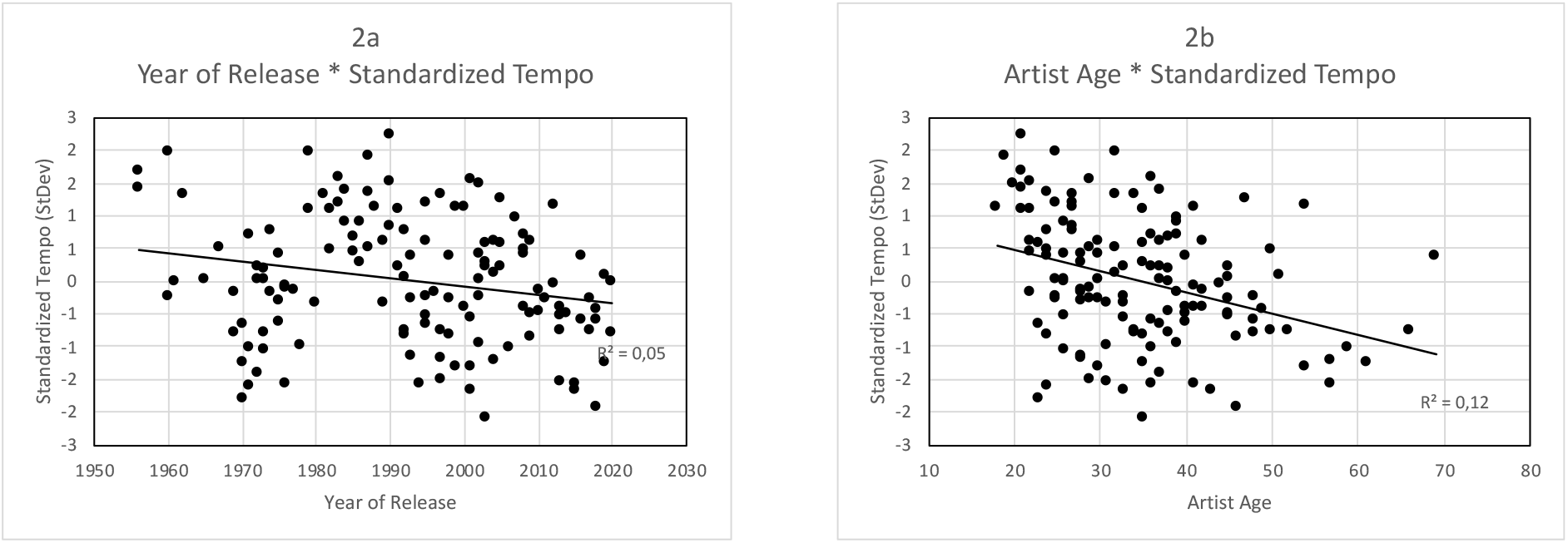
Standardised Tempo as a function of a) Year of Release and b) Artist Age.

### Between-artist variation

Figure 3 a-j shows *Standardized Tempo* plotted against *Age* separately for each artist. A series of simple OLS linear regressions revealed that all 10 artists exhibited a downward trend in *Standardized Tempo* as they increased in age, supporting *H1*. This downward trend is apparent regardless of the time period (e.g., decade(s)) during which an artist released their music. Moreover, with data plotted individually for each artist, noticeable between-artist variation in evolution of tempo across time becomes apparent, supporting *H2*. Five of these trends were statistically significant: Elvis Presley, *F*(1, 19) = 15.84, *R*^*2*^ = .46, *p* < .001, Eminem, *F*(1, 8) = 18.44, *R*^*2*^ = .70, *p* = .003, Lil Wayne, *F*(1, 9) = 11.39, *R*^*2*^ = .56, *p* = .008, Michael Jackson, *F*(1, 4) = 39.72, *R*^*2*^ = .91, *p* = .003, and Whitney Houston, *F*(1, 4) = 11.42, *R*^*2*^ = .74, *p* < .05. Overall, the amount of variance in tempo explained by age ranged from 1% (Elton John) to 91% (Michael Jackson).

**Figure 3.**
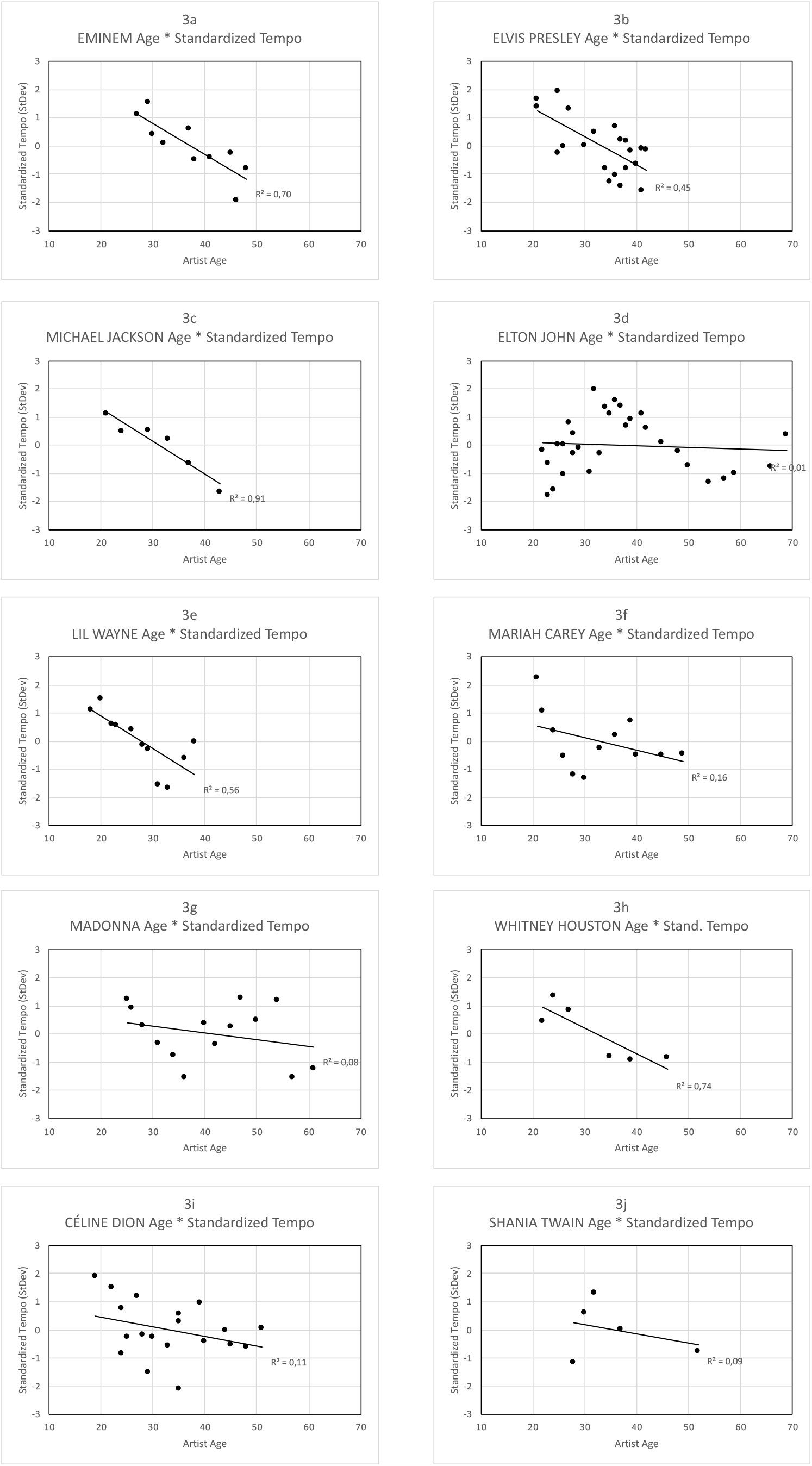
Standardised Tempo as a function of Age for a) Eminem, b) Elvis Presley, c) Michael Jackson, d) Elton John, e) Lil Wayne, f) Mariah Carey, g) Madonna, h) Whitney Houston, i) Céline Dion, and j) Shania Twain.

### Chronological age as a predictor of musical tempo

To investigate if artist age predicts tempo of musical output more generally, *Standardized Tempo* was averaged by age across all artists (*Averaged Standardized Tempo*). To limit the impact of individual albums or artists, values based on fewer than two albums by different artists were excluded (see Figure 4). *Age* was subsequently entered into a simple OLS linear regression as a predictor of *Averaged Standardized Tempo*. Results revealed a robust effect of *Age, F*(1, 26) = 10.49, *R*^*2*^ = 0.29, *p* = .003, supporting *H3*. Overall, musical tempo was observed to decrease by almost one and a half standard deviations from artists’ early twenties to their late fifties. In other words, regardless of the calendar year in which an artist released an album, their output tended to decrease in tempo as they aged.

**Figure 4.**
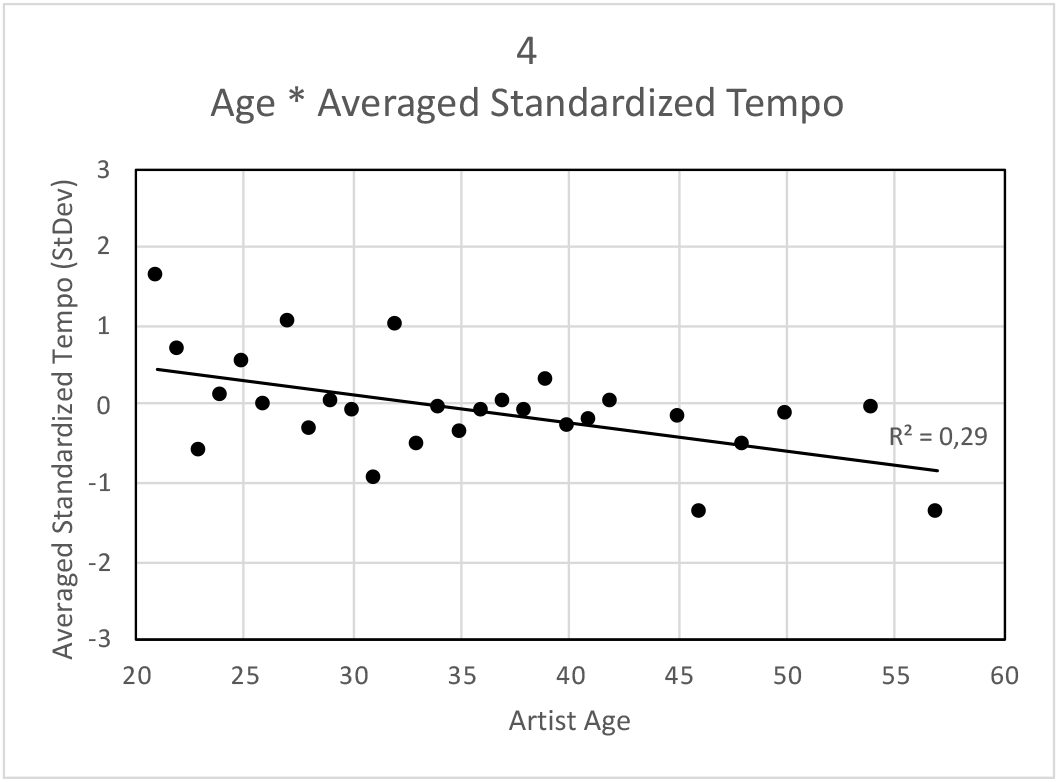
Averaged Standardised Tempo as a function of Artist Age.

## Discussion

The present study tested 3 hypotheses: (*H1*) That recording artists would exhibit a downward trend in tempo of their output across their careers, driven by the intimate connection between music performance and biological motion and a well-documented slowdown in motor tempo across the adult lifespan; (*H2*) that artist-specific variations would be observed in these trends due to individual differences in motor tempo and impacts arising from different producers, musicians, label executives, and genres; and (*H3*) that by nesting tempo within-artist it would be possible to construct a robust overall model in which artist age predicts tempo of musical output. All three hypotheses were confirmed. Age was found to explain up to 91% of the variance in tempo of individual artists’ musical output. Across all artists, tempo was observed to decrease by close to one and a half standard deviations from their early twenties to their late fifties.

Despite significant individual variation (*H2*), therefore, chronological age predicted the tempo of music released by the world’s most successful artists of the popular music era (*H1, H2*). That such a clear effect of age was identified despite influences from different personnel as well as across genres, makes the effect all the more striking. Furthermore, these results demonstrate that commercial recordings, millions of which are instantly accessible via a range of streaming platforms, can offer profound insights into a fundamental and understudied aspect of human functioning across the lifespan.

These results have implications concerning the extent to which artists (are able to) both shape their compositions and engage their audience. By definition, physical constraints on motor speed influence the frequency of events an individual can comfortably produce or would prefer to execute.^48^ From a musician’s perspective, biophysiological constraints on performance-critical motor actions, such as rhythmic manipulations of the vocal tract, breathing apparatus, upper body, arms, and fingers, will thus shape their rhythmic capacities. This includes the range of musical tempi they prefer or are able to produce. Because of this, as musicians age, their ability to express particular emotions, adhere to genre or stylistic characteristics, or create and release tension in their audience could be compromised. This could transform the characteristics and character of their musical output, in turn modifying how listeners respond. In particular, changes in musical features will affect listeners’ perception of expression, their level of physiological arousal, and characteristics of their music-induced body movement. All of which will be mediated by the degree to which an artist’s performance is connected to their own physicality.

Additional support for attributing the decrease in artists’ tempo observed in the present study to a degradation in their motor capacities can be found in notable characteristics exhibited by four of the five artists for whom age significantly predicted tempo. Both Elvis Presley and Michael Jackson routinely danced to their music when performing. In fact, both were known as much for their body movement and dance styles as for their music. For these two artists, body movement and music were virtually inseparable. The underlying emphasis on physicality and rhythmicity in the vocal delivery of rap lyrics^49^ also connects both Eminem and Lil Wayne more viscerally to their music than artists performing in other styles. For these four artists, body movement and music are or were intimately intertwined. Consequently, as they (have) aged, the gradual degradation of their motor capacities (has) had a more pronounced impact on the tempo of their musical output.

Overall, the results are consistent with the slowing-with-age hypothesis, which proposes a slowdown in behavioural speed across the adult lifespan. This slowdown is believed to affect motor competence, especially motor speed, due to a reduction in the limits of processing speed and neuromuscular changes including reduced muscle strength and endurance.^50^ Notably, networks including the prefrontal cortex and basal ganglia are typically more implicated in motor control later in life, brain regions frequently impaired in older age.^51^ This last point raises an additional implication of the present study: The potential for neurocognitive decline to be accelerated by typical characteristics of musicians’ lifestyles and personality.

Musicians are more likely to abuse alcohol and drugs^52^ and display higher levels of stress and anxiety.^53^ Musicians also maintain more irregular schedules and exhibit a higher prevalence of mental and physical health issues compared to the general population.^54^ The pronounced effects seen in the present study might indicate that such behaviours and characteristics accelerate decline in basic motor speed – and thus musical tempo – across the lifespan. It seems notable that three of the five artists whose musical tempo declined statistically significantly over their careers died prematurely from drug and alcohol abuse.

This raises the tantalizing possibility of using musical tempo to predict an artist’s physical or mental wellbeing. Indicators of slower SMT, such as walking pace, are in fact related to increased incidence of critical health issues including stroke.^55^ Walking pace has also been shown to indicate premature ageing and predict associated risk factors such as decline in both neurocognitive^56^ and physical function.^57^ A lot more work is clearly needed to tease apart these various effects and influences, but the connections here are striking.

It would also seem pertinent to consider not only effects of physical ageing, but those resulting from psychological maturation as well. Younger people tend to engage with music for identity, positive and negative mood management, reminiscence, diversion, arousal, and social interaction,^58^ while older individuals do so for entertainment, connection, wellbeing, time management, therapeutic benefits and spirituality.^59^ A recent study suggests that music preferred by older individuals contains higher levels of love-tenderness and lower levels of pain-sadness compared with music listened to by younger people.^60^ As a consequence, different reasons for engaging with music will likely lead to differences in audio characteristics of the music one engages with.^61^ To what extent this applies to artists and the music they create at different stages of the life course remains an open question, but one that deserves further investigation.

Finally, while this work offers support for a connection between age and tempo of artistic output, it remains unclear how generalizable this connection is beyond a sample of the most culturally significant artists. To gain a comprehensive picture of this potentially significant phenomenon, future work should investigate a broader range of musicians, musical styles, and musical periods.

## Conclusions

Timing is a critical aspect of musical performance intimately related to the dynamics of biological motion. The aim in the present study was to investigate the impact of ageing and concomitant degradation in motor speed on this relationship. Analysis of a large corpus of culturally-significant and genre-diverse music recordings revealed a robust negative relationship between age and physical tempo independent of broader musical trends. The impact of age on tempo was overwhelming for artists most physically connected with their music. Given the ubiquity of music across human cultures and the powerful effects it exerts on listeners’ emotional and psychobiological functioning, these results have important implications concerning the degree to which artists (are able to) both shape their compositions and engage their audience as they age.

## References

1 T. Hedden, J. D. Gabrieli. Insights into the ageing mind: A view from cognitive neuroscience. Nat. Rev. Neurosci. 5(2), 87–96 (2004).

2 D. C. Park, P. Reuter-Lorenz. The adaptive brain: Aging and neurocognitive scaffolding. Annu. Rev. Psychol. 60, 173–196 (2009).

3 I. J. Deary, J. Corley, A. J. Gow, S. E. Harris, L. M. Houlihan, R. E. Marioni, L. Penke, S. B. Rafnsson, J. M. Starr. Age-associated cognitive decline. Br. Med. Bull. 92(1), 135–152 2009.

4 G. Kavé, A. Knafo, A. Gilboa. The rise and fall of word retrieval across the lifespan. Psychol. Aging 25(3), 719 (2010).

5 F. Z. Esfahlani, J. Faskowitz, J. Slack, B. Mišić, R. F. Betzel. Local structure-function relationships in human brain networks across the lifespan. Nat Commun. 13, 2053 2022.

6 A. Baudouin, S. Vanneste, M. Isingrini. Age-related cognitive slowing: The role of spontaneous motor tempo and processing speed. Exp. Aging Res. 30:3, 225–239 (2004).

7 S. K. Hunter, M. W. Thompson, R. D. Adams. Reaction time, strength, and physical activity in women aged 20–89 years. J. Aging Phys. Act. 9(1), 32–42 (2001).

8 M. Morgan, J. G. Phillips, J. L. Bradshaw, J. B. Mattingley, R. Iansek, J. A. Bradshaw. Age-related motor slowness: Simply strategic? J. Gerontol. 49(3), M133–M139 (1994).

9 T. A. Salthouse. The processing-speed theory of adult age differences in cognition. Psychol. Rev. 103(3), 403–428 (1996).

10 C. Voelcker-Rehage. Motor-skill learning in older adults—a review of studies on age-related differences. Eur. Rev. Aging Phys. Act. 5(1), 5–16 (2008).

11 M. Broughton, C. Stevens. Music, movement and marimba: An investigation of the role of movement and gesture in communicating musical expression to an audience. Psychol. Music 37(2), 137–153 (2009).

12 J. W. Davidson. Visual perception of performance manner in the movements of solo musicians. Psychol. Music 21(2), 103–113 (1993).

13 J. W. Davidson. Qualitative insights into the use of expressive body movement in solo piano performance: a case study approach. Psychol. Music 35(3), 381–401 (2007).

14 J. Juchniewicz. The influence of physical movement on the perception of musical performance. Psychol. Music 36(4), 417–427 (2008).

15 M. M Wanderley, B. W. Vines, N. Middleton, C McKay, W. Hatch. The musical significance of clarinetists’ ancillary gestures: An exploration of the field. J. New Music Res. 34(1), 97–113 (2005).

16 A. Friberg, J. Sundberg. Does music performance allude to locomotion? A model of final ritardandi derived from measurements of stopping runners. J. Acoust. Soc. Am. 105(3), 1469–1484 (1999).

17 C. Palmer. Sequence memory in music performance. Curr. Dir. Psychol. Sci. 14(5), 247–250 (2005).

18 C. Palmer, P. Q. Pfordresher. Incremental planning in sequence production. Psychol. Rev. 110(4), 683–712 (2003).

19 F. T. Van Vugt, H-C. Jabusch, E. Altenmüller. Fingers phrase music differently: trial-to-trial variability in piano scale playing and auditory perception reveal motor chunking. Front. Psychol. 3(495) (2012).

20 B. Sievers, L. Polansky, M. Casey, T. Wheatley. Music and movement share a dynamic structure that supports universal expressions of emotion. Proc. Natl. Acad. Sci. U.S.A. 110(1), 70–75 (2013).

21 S. Arom, M. Thom, B. Tuckett, R. Boyd. African polyphony and polyrhythm: Musical structure and methodology. Cambridge, UK: Cambridge University Press (2004).

22 I. Cross. Music, cognition, culture, and evolution. Ann. N. Y. Acad. Sci. 930(1), 28–42 (2001).

23 N. L. Wallin, B. Merker, S. Brown (Eds.). The origins of music. Cambridge, MA: MIT Press (2000).

24 V. S. Mattay, F. Fera, A. Tessitore, A. R. Hariri, S. Das, J. H. Callicott, D. R. Weiberger. Neurophysiological correlates of age-related changes in human motor function. Neurology 58(4), 630–635 (2002).

25 T. A. Salthouse. The processing-speed theory of adult age differences in cognition. Psychol. Rev. 103(3), 403–428 (1996).

26 P. Verhaeghen, D. W. Steitz, M. J. Sliwinski, J. Cerella. Aging and dual-task performance: A meta-analysis. Psychol. Aging 18(3), 443–460 (2003).

27 T. D. Lee, L. R. Wishart, J. E. Murdoch. Aging, attention, and bimanual coordination. Can. J. Aging 21(4), 549–557 (2002).

28 S. Vanneste, V. Pouthas, J. H. Wearden. Temporal control of rhythmic performance: A comparison between young and old adults. Exp. Aging Res. 27(1), 83–102 (2001).

29 A. Baudouin, S. Vanneste, M. Isingrini. Age-related cognitive slowing: The role of spontaneous motor tempo and processing speed. Exp. Aging Res. 30(3), 225–239 (2004).

30 D. Hammerschmidt, K. Frieler, C. Wöllner. Spontaneous motor tempo: Investigating psychological, chronobiological, and demographic factors in a large-scale online tapping experiment. Front. Psychol. 12:677201 (2021).

31 J. D. McAuley, M. R. Jones, S. Holub, H. M. Johnston, N. S. Miller. The time of our lives: Life span development of timing and event tracking. J. Exp. Pschol. Gen. 135(3), 348–367 (2006).

32 K. Hine, K. Abe, Y. Kinzuka, M. Shehata, K. Hatano, T. Matsui, S. Nagauchi. Spontaneous motor tempo contributes to preferred music tempo regardless of music familiarity. Front. Psychol. 13:952488 (2022).

33 N. L. Wallin, B. Merker, S. Brown (Eds.). The origins of music. Cambridge, MA: MIT Press (2000).

34 H. Sigmundsson, H. Lorås, M. Haga. Assessment of motor competence across the life span: Aspects of reliability and validity of a new test battery. SAGE Open 6(1) (2016).

35 T. Eerola, J. K. Vuoskoski. A review of music and emotion studies: Approaches, emotions, models, and stimuli. Music Percept. 30(3), 307–340 (2013).

36 T. LH. Li, A. B. Chan. Genre classification and the invariance of MFCC features to Key and Tempo. In International Conference on MultiMedia Modeling (pp. 317–217). Berlin, Heidelberg; Springer (2011).

37 M. Goodchild, B. Gingras, S. McAdams. Analysis, performance, and tension perception of an unmeasured prelude for harpsichord. Music Percept. 34(1), 1–20 (2016).

38 G. D. Webster, C. G. Weir. Emotional responses to music: Interactive effects of mode, texture, and tempo. Motiv. Emot. 29(1), 19–39 (2005).

39 L. O. Lundqvist, F. Carlsson, P. Hilmersson, P. N. Juslin. Emotional responses to music: Experience, expression, and physiology. Psychol. Music 37(1), 61–90 (2009).

40 B. Burger, M. R. Thompson, G. Luck, S. Saarikallio, P. Toiviainen. Hunting for the beat in the body: On period and phase locking in music-induced movement. Front. Hum. Neurosci. 8(903) (2014).

41 https://www.riaa.com

42 C. Drake, L. Gros, A. Penel. How fast is that music? The relation between physical and perceived tempo. In S. W. Yi (Ed.), Music, mind, and science (pp. 190–203), Seoul, Korea: Seoul National University (1999).

43 C. Sachs. Rhythm and Tempo. London: Dent. 1953.

44 A. Baudouin, S. Vanneste, M. Isingrini. Age-related cognitive slowing: The role of spontaneous motor tempo and processing speed. Exp. Aging Res. 30(3), 225–239 (2004).

45 J. D. McAuley, M. R. Jones, S. Holub, H. M. Johnston, N. S. Miller. The time of our lives: Life span development of timing and event tracking. J. Exp. Pschol. Gen. 135(3), 348–367 (2006).

46 D. Rose, D. J. Cameron, P. J. Lovatt, J. A. Grahn, L. E. Annett. Comparison of spontaneous motor tempo during finger tapping, toe tapping, and stepping on the spot in people with and without Parkinson’s disease. J. Mov. Disord. 13(1), 47–56 (2020).

47 S. Vanneste, V. Pouthas, J. H. Wearden. Temporal control of rhythmic performance: A comparison between young and old adults. Exp. Aging Res. 27(1), 83–102 (2001).

48 P. Brown. An enquiry into the origins and nature of tempo behaviour. Psychol. Music 7(1), 19–35 (1979).

49 A. Krims. Rap Music and the Poetics of Identity. Cambridge, UK: Cambridge University Press (2000).

50 A. Baudouin, S. Vanneste, M. Isingrini. Age-related cognitive slowing: The role of spontaneous motor tempo and processing speed. Exp. Aging Res. 30(3), 225–239 (2004).

51 R. D. Seidler, J. A. Bernard, T. B. Burutolu, B. W. Fling, M. T. Gordon, J. T. Gwin, Y. Kwak, D. B. Lipps. Motor control and aging: Links to age-related brain structural, functional, and biochemical effects. Neurosci. Biobehav. Rev. 34(5), 721–733 (2010).

52 K. E. Miller, B. M. Quigley. Sensation-seeking, performance genres and substance use among musicians. Psychol. Music, 40(4), 389–410 (2011).

53 A. Détári, H. Egermann, O. Bjerkeset, J. Vaag. Psychosocial work environment among musicians and in the general workforce in Norway. Front. Psychol. Front. Psychol. 11:1315. doi: 10.3389/fpsyg.2020.01315

54 G. Musgrave. Music and wellbeing vs. musicians’ wellbeing: examining the paradox of music-making positively impacting wellbeing, but musicians suffering from poor mental health. Cult. Trends, 1–16 (2022).

55 S. Hayes, J. F. Forbes, C. Celis-Morales, J. Anderson, L. Ferguson, J. M. Gill, … J. Pell. Association Between Walking Pace and Stroke Incidence: Findings from the UK Biobank Prospective Cohort Study. Stroke 51(5), 1388–1395 (2020).

56 R. A. Hackett, H. Davies-Kershaw, D. Cader, M. Orrell, A. Steptoe. Walking speed, cognitive function, and dementia risk in the English longitudinal study of ageing. J. Am. Geriatr. Soc. 66, 1670–1675 (2018).

57 L. J. H. Rasmussen, A. Caspi, A. Ambler, J. M. Broadbent, H. J. Cohen, T. d’Arbeloff, T., … T. E. Moffitt. Association of neurocognitive and physical function with gait speed in midlife. JAMA Netw. Open 2(10), e1913123–e1913123 (2019).

58 A. J. Lonsdale, A. C. North. Why do we listen to music? A uses and gratifications analysis. Br. J. Psychol. 102(1), 108–134 (2011).

59 T. Hays, V. Minichiello. The contribution of music to quality of life in older people: An Australian qualitative study. Ageing Soc. 25(2), 261–278 (2005).

60 A. Mavrolampados, D. Duman, I. Burunat, N. Snape, P. Neto, M. Hartmann, …, P. Toiviainen, (in prep). Impact of age and gender on affective experiences in music.

61 D. Duman, P. Neto, A. Mavrolampados, P. Toiviainen, G. Luck. Music we move to: Spotify audio features and reasons for listening. Plos one 17(9), e0275228 (2022).

